# Antibiotic-producing Micrococcales govern the microbiome that inhabits the fur of two- and three-toed sloths

**DOI:** 10.1101/2022.04.08.486316

**Authors:** Diego Rojas-Gätjens, Katherine S. Valverde-Madrigal, Keilor Rojas-Jimenez, Reinaldo Pereira, Judy Avey-Arroyo, Max Chavarría

**Author notes:** **Correspondence to: Max Chavarría**, Escuela de Química & Centro de Investigaciones en Productos Naturales (CIPRONA), Universidad de Costa Rica, Sede Central, San Pedro de Montes de Oca, San José, 11501-2060, Costa Rica.

## Abstract

Sloths have a dense coat on which insects, algae, and fungi coexist in a symbiotic relationship. This complex ecosystem requires different levels of control, however, most of these mechanisms remain unknown. We investigated the bacterial communities inhabiting the hair of two- (*Choloepus Hoffmani*) and three-toed (*Bradypus variegatus*) sloths and evaluated their potential for producing antibiotic molecules capable of exerting control over the hair microbiota. The analysis of 16S rRNA amplicon sequence variants (ASVs) revealed that the communities in both host species are dominated by Actinobacteriota and Firmicutes. The most abundant genera were *Brevibacterium, Kocuria/Rothia, Staphylococcus, Rubrobacter, Nesterenkonia, and Janibacter*. In addition, we isolated nine strains of *Brevibacterium* and *Rothia* able to produce substances that inhibited the growth of common mammalian pathogens. The analysis of the biosynthetic gene clusters (BCGs) of these nine isolates suggests that the pathogen-inhibitory activity could be mediated by the presence of siderophores, terpenes, beta-lactones, Type III polyketide synthases (T3PKS), ribosomally synthesized, and post-translationally modified peptides (RiPPs), non-alpha poly-amino acids (NAPAA) like e-Polylysin, ectoine or nonribosomal peptides (NRPs). Our data suggest that Micrococcales inhabiting sloth hair could have a role in controlling microbial populations in that habitat, improving our understanding of this highly complex ecosystem.

## Introduction

Almost all mammals have fur or other hair-related structures that help them adapt to their habitats. Hair usually provides thermal regulation and camouflage (Dawson et al., 2014) also it has been related to some communication and mating processes (Buffoli et al., 2014). Beyond their functional role for animals, hair constitutes a habitat for multiple microorganisms on which they coexist and associate in complex ways with their host (Chen et al., 2018). In mammals, due to their anatomical proximity, it is difficult to separate the microbiome associated with skin and fur, and the vast majority of studies refer to the skin microbiome (Belheouane et al., 2020; Chen et al., 2018; Council et al., 2016; Ross et al., 2019). For example, a high microbial diversity has been reported in human skin (Byrd et al., 2018), and it has been shown that it presents a high proportion of symbionts that control the population of pathogenic bacteria through the secretion of multiple compounds such as bacteriocins or fatty acids (Byrd et al., 2018; O’Sullivan et al., 2019). For its part, the hair microbiota in humans has been shown to have a similar composition to skin samples (Polak-Witka et al., 2020; Saxena et al., 2021). The most abundant genera found in hair follicles correspond to *Cutibacterium acnes* (previously known as *Propionibacterium acnes*), *Staphylococcus* sp., *Corynebacterium, Streptococcus, Acinetobacter, Prevotella* and *Malassezia* (Lousada et al., 2021; Polak-Witka et al., 2020; Watanabe et al., 2021), however, it is important to mention that this microbiota is influenced by multiple factors including demography, use of cosmetics products or oils and genetics (Brinkac et al., 2018).

The human skin (and fur) microbiome is the most studied; the microbiome present in the skin and hair of other mammals and their possible functions are poorly understood or remain completely unknown. Understanding these microbe-mammalian hair relationships is important not only from an ecological perspective but also because we could reveal microorganisms, enzymes or small molecules of biotechnological interest. A global analysis of multiple animals’ skin microbiome, based on 16S rRNA amplicon sequencing revealed that the skin of most mammals differs significantly from the human skin microbiome and resembles soil microbiome in most cases, which is probably a consequence of the constant environmental exposure of their fur. Also, a phylosymbiosis of the microbiota was found among the different mammals’ clades. Bacteria belonging to *Staphylococcus, Corynebacterium*, and *Micrococcus* genera are reported as recurrent members on the skin and fur of mammals (Kloss et al., 1976; Ross et al., 2019). In non-human primates, it was revealed that even though members of the *Staphylococcus* and *Corynebacterium* genera were present, other phylotypes such as *Anaerococcus* and *Prevotella* are an abundant part of the microbial community (Council et al., 2016). Furthermore, on livestock animals, it has been shown that the skin microbiome has an important prevalence of Bacteoidetes, Chloroflexi, and Proteobacteria (Ross et al., 2018; Ross et al., 2019). The role of these phylotypes in the animal’s physiology is unclear. Nevertheless, it has been postulated they can be involved in protection from pathogenic bacteria via competition or maturation of the immune system (Bradley et al., 2016; Rodrigues Hoffman, 2017).

Sloths are mammals whose skin and hair microbiota is partially known (Higginbotham et al., 2014; Pauli et al., 2014). Sloths are mammals adapted to live in the canopy layer of the rainforest. These animals are characterized as being solitary with parsimonious movements and a relatively low metabolic rate compared to other mammals (Falconi et al., 2015; Taube et al., 2008). They have a low body temperature, making them intolerant to sudden variations in temperature; for this reason, they spend most of their time sunbathing and feeding (De Oliveira et al., 2014). There are a total of six different sloth species divided into two genera: *Bradypus* (two-toed sloths), with four species, and *Choloepus* (three-toed sloths), with two species (Dill-McFarland et al., 2016; Mendoza et al., 2015). In Costa Rica, only two species are present *B. variegatus* and *C. hoffmanni* (Vaughan et al., 2007). These two species were declared in August 2021 as a national symbol in the country due to their importance in touristic activities and their ecological roles (see Figure 1).

**Figure 1.**
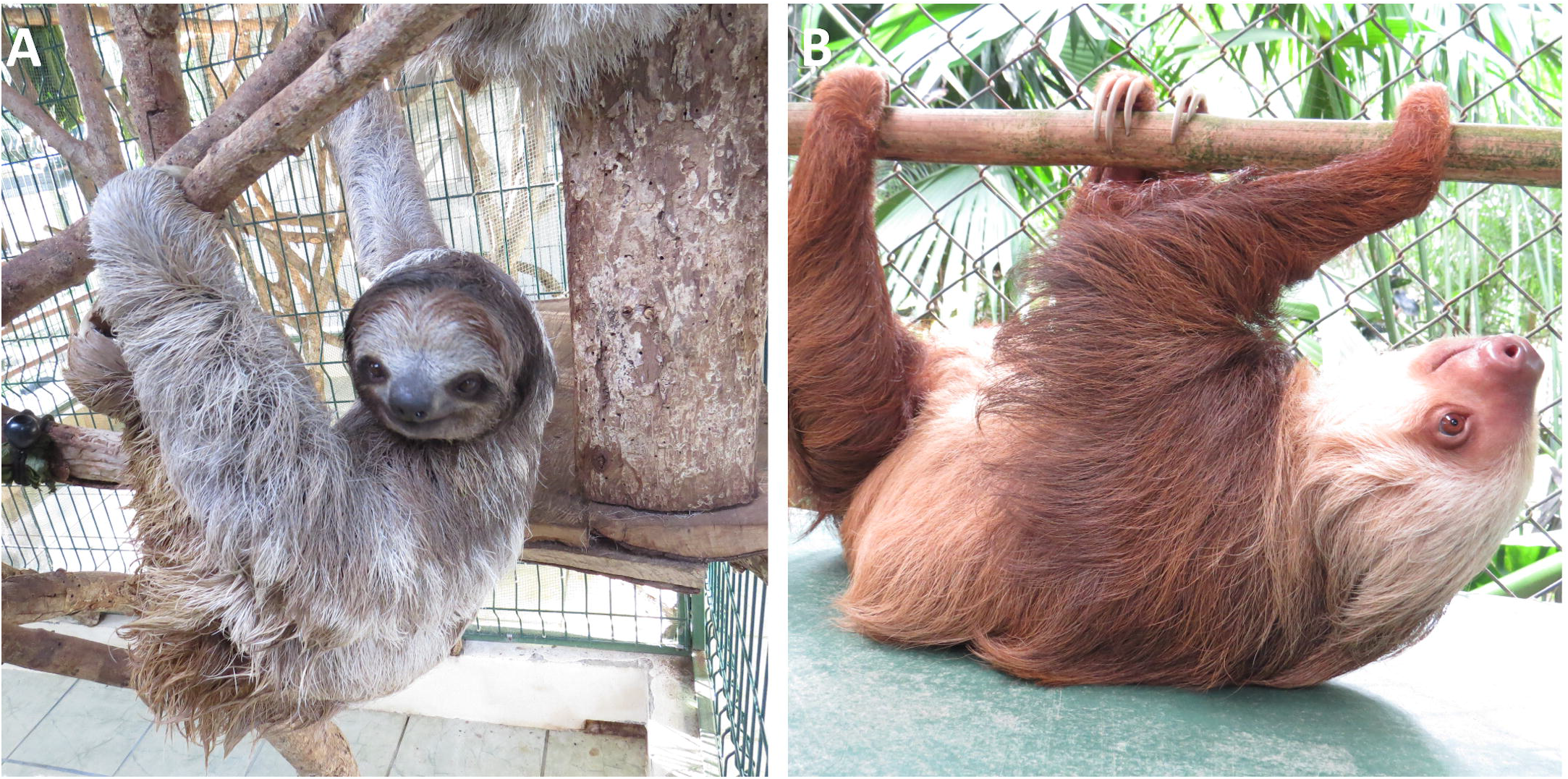
Sloths that inhabit in the Sloth Sanctuary of Costa Rica located in Cahuita, Costa Rica. A) *Choloepus hoffmani*, B) *Bradypus variegatus*.

Sloth fur is a dense coat on which macro- and microorganisms coexist (see Figure 1). It is usually covered with green algae from the genus *Trichophilus*, which helps sloths camouflage from predators (Suutari et al., 2010). Also, a moth classified as *Cryptoses* lives on sloths’ hair and is in charge of the nitrification process that supports the growth of *Trichophilus* (Waager and Montgomery, 1976). Previously, a study that focused on culturable fungi that inhabit two-toed sloth’s hair revealed that these fungi secrete multiple compounds that inhibit the growth of numerous bacteria and parasites such as *T. cruzi* (Higginbotham et al., 2014). Even though sloths fur has been proved to be a complex habitat with various organisms occupying it, little information exists about its bacterial community, functions, and possible roles as symbionts of these mammals. In this work, we characterize the bacterial community inhabiting the hair of both sloth species from Costa Rica (*Choloepus hoffmani* and *Bradypus variegatus*) (see Figure 1). Additionally, we intend to explore the possible roles of these bacteria in the pathogen control of the microbial community on sloth hair.

## Experimental Procedures

### Sampling and permits

All samples of sloth hair were obtained from the Sloth Sanctuary (http://www.slothsanctuary.com) in Cahuita, Limon (9°47’58.5”N 82°54’53.1”W) in February 2020. The sanctuary shelters around 250 individuals of *Bradypus variegatus* and *Choloepus hoffmani* (see Figure 1). The Sloth Sanctuary has been operating since 1992 and is dedicated to the rescue, rehabilitation, and research of sloths and conserving their rainforest habitat. All the sloths have come to the sanctuary because they were abandoned by their mothers, or have suffered some kind of accident. In cases where it is possible, the animals are reinserted into their natural habitats, however, many do not have that capacity and are therefore kept in the refuge. All sloths are kept in cages which can be either alone or with partners of the same species. In the case of shared cages, only one sloth per cage was sampled. Hair samples were taken from the back of the sloth usinors and then stored in sterile 50 mL conical tubes. We obtained two subsamples from 13 individuals of *Bradypus* and 15 individuals of *Choloepus*. The names, code, years in the refuge, and other related characteristics of each sampled animal are shown in Table S1. Samples for total DNA extraction were stored at -20 °C until processed. The samples for bacterial isolation were immediately transported to the laboratory and processed as described below. Permits to sample were obtained from the Institutional Commission of Biodiversity of the University of Costa Rica (resolution Nº 253) and Judy Avey-Arroyo, the property owner.

### Scanning electron microscopy

Hairs of *B. variegatus* and *C. hoffmani* ∼5 mm long were fixed with osmium tetroxide vapor (4%, in 0.2 M sodium phosphate buffer, pH 7.2). The hair fragments were placed in 2-cm-diameter plastic lids, which were kept inside a 15-cm-diameter glass Petri dish containing filter paper moistened with the osmium solution for 48 hours in a hood of extraction. After this time, the material was removed from the Petri dish and left to dry in the fume hood for 12 hours. The hairs were placed on an aluminum base containing double-sided carbon conductive tape and plated with gold in a Denton Vaccum Desk V ion spreader (20mA, 3 minutes). Subsequently, the hairs were observed in scanning electron microscopy (JEOL JSM 6390LV) at 20 KV.

### Total DNA extraction and sequencing

For total DNA extraction of sloth hair, the samples were proceeded as described by Rojas-Gätjens (2021) with some modifications. Briefly, hair samples were cut with sterile scissors into smaller pieces. A total of 300-500 mg of hair per sample was extracted with a DNA isolation kit (PowerSoil®, MoBio, Carlsbad, CA, USA) as described by the manufacturer. Cell lysis was accomplished by two steps of bead beating (FastPrep-24, MP Biomedicals, Santa Ana, CA, USA) for 30 s at 6.0 m s^−1^. For the construction of the microbial amplicon library (16S rRNA), the V4 hypervariable region was PCR-amplified with universal primers 515F (5’-GTGCCAGCMGCCGCGGTAA-3’) and 806R (5’-GGACTACHVGGGTWTCTAAT-3’) (Carporaso et al., 2010). The PCR-generated products were subjected to a 250 nt paired-end sequencing (Novoseq, Novogene).

### Bioinformatic and statistical analysis

Amplicon sequencing data was analyzed using the DADA2 pipeline (version 1.12), as described by Rojas-Gätjens et al. (2021). Briefly, reads were quality-filtered, deduplicated, and amplicon sequence variant (ASV), a higher-resolution analog of the traditional operational taxonomic unit (OTU), was inferred. The chimeric sequences were removed. This process resulted in an ASV table, which records the number of times each exact amplicon sequence variant was observed in a sample. ASVs were taxonomically assigned by comparing sequences against the SILVA reference database v138 (Quast et al., 2013) and then curated by comparing sequences against the GenBank database of NCBI. Global singletons, doubletons, and sequences classified as Eukarya, Chloroplast, and Mithocondria were removed. The average number of sequences per sample obtained was 132265, ranging from 91415 to 170677. For estimation of the core microbiome of both sloths, we use the package R package microbiome (Lahti and Shetty, 2018). The core microbiome was defined with a prevalence of 50% and abundance of 1% according to Lahti and Shetty (2018). The raw sequencing data were deposited in the sequence-read archive (SRA) of Genbank under the Bioproject ID PRJNA804868

The statistical analysis and their visualization were performed with R statistical program (R-core-Team 2019) and the Project Jupyter notebook interface. The package Vegan v2.5-6 (Oksanen et al. 2019) was used to calculate alpha diversity estimators (Observed abundance, Shannon index, and Simpson index) and Non-metric multidimensional scaling analysis (NMDS). Wilcoxon test was performed for comparison of the alpha diversity estimators between hosts (*Choloepus hoffmani* and *Bradypus variegatus*). Data tables with the ASV abundances were normalized into relative abundances and then converted into a Bray–Curtis similarity matrix. The effects of the hosts and sex on the bacterial community composition were evaluated with non-parametric multivariate analysis of variance (PERMANOVA) and pairwise PERMANOVA (adonis2 function with 999 permutations). The effect of the time they present in captivity was evaluated using the envfit function developed on the package Vegan v2.5-6 (Oksanen et al. 2019).

### Isolation of antibiotic-producing bacteria

Samples were processed as described by Cambronero-Heinrichs (2019). Briefly, samples were diluted in PBS 1X (Na_2_HPO_4_ 0.01 mol L^-1^, KH_2_PO_4_ 0.0018 mol L^-1^, NaCl 0.137 mol L^-1^, KCl 0.0027 mol L^-1^) employing a 1:1 sample weight/volume ratio and then serially diluted until 10^−3^. Dilutions were pre-incubated (60°C for 10 min) for avoiding the growth of Enterobacteria. Dilutions were plated on oatmeal agar (Jensen et al., 2005) supplemented with nalidixic acid (25 mg ml^−1^) and cycloheximide/clotrimazole (50 mg ml^−1^). Plates were incubated for three months (30°C) and examined every week. All bacteria were transferred to ISP2 medium (Pridham et al., 1957). All isolates were preserved (−80°C, 15 % glycerol).

### Screening of antimicrobial activity

The ability of isolates to inhibit bacterial growth was evaluated by the agar disc diffusion method described by Cambronero-Heinrichs (2019). Since there are no known possible pathogens on sloth skin, we decided to perform antibiotic activity tests against ATCC model microorganisms such: *Escherichia coli* ATCC25922, *Pseudomonas aeruginosa* PAO1, *Staphylococcus aureus* ATCC25923, *Candida albicans* ATCC10231, and *Bacillus subtilis subsp. spizizenii* ATCC6336. In addition, we included in the test set other available microorganisms known to be pathogenic in mammals (i.e. *Streptococcus agalactiae, Staphylococcus epidermidis, Staphylococcus saprophyticus, Kleibsella pneumonie, Serratia marcescens)*, as well as some genera of fungi that were isolated from the hair of sloths previously (i.e. *Cytospora sp*., *Fusarium sp*., *Colletrichum sp*., *Hypocrea sp*., *Trichoderma sp*) (Higginbotham et al., 2014).

Bacteria and yeast pathogens strains were individually inoculated in either Luria–Bertani agar or potato dextrose agar (OD = 0.1, from overnight cultures). A piece of agar (diameter 6mm) with confluent growth of each actinobacterial isolate (2-week-old cultures in ISP2) was cut and placed on the surface of the medium, with bacterial growth uppermost. A Whatman® cellulose filter paper inoculated with kanamycin (5 uL, 5 mg/mL; 50 mg/mL for *P. aeruginosa*) or clotrimazole (5 mg/mL) was used as a positive control. A filter without any substance was used as the negative control. The plates were incubated for 24h at 37 °C for *E. coli, P. aeruginosa, S. aureus, Streptococcus agalactiae, Staphylococcus epidermidis, Staphylococcus saprophyticus, Kleibsella pneumonie, Serratia marcescens*, and *C. albicans*, and 30° C for all others. Every experiment was performed in duplicate. Inhibition halos were measured in centimeters, and the results were ranked by size (see Table 1).

**Table 1.** Identity and antimicrobial activity of bacteria isolated from sloths’ hair.

| Strain | Host | Activity against pathogen |  |  |  |  |  |  |  |  |  |  |  |  |  |  |
| --- | --- | --- | --- | --- | --- | --- | --- | --- | --- | --- | --- | --- | --- | --- | --- | --- |
|  |  | SA | EC | PA | BS | CA | SAg | SS | SE | KP | SM | Cy | Co | Fu | Hy | Tr |
| *Rothia koreensis* 10C1 | *Choloepus* | ++ | - | - | - | - | - | - | - | - | - | - | - | - | - | - |
| *Rothia kristinae* 13C9 | *Choloepus* | - | - | + | - | - | - | - | - | - | - | + | - | - | - | - |
| *Rothia kristinae* 13C8 | *Choloepus* | + | - | + | +++ | - | - | - | - | - | - | + | - | - | - | - |
| *Rothia* sp. 9C1 | *Choloepus* | + | - | + | +++ | - | - | - | - | - | - | - | - | - | - | - |
| *Rothia kristinae* 11C2 | *Choloepus* | + | - | + | + | - | - | - | - | - | - | + | - | - | - | - |
| *Rothia kristinae* 6B2 | *Bradypus* | - | ++ | ++ | - | - | - | - | - | - | - | - | - | - | - | - |
| *Rothia kristinae* 3B2 | *Bradypus* | + | - | + | +++ | - | + | - | - | - | - | - | - | - | - | - |
| *Brevibacterium sediminis*<br>11B2 | *Bradypus* | - | ++ | - | - | - | - | - | - | - | - | - | - | - | - | - |
| *Rothia kristinae* 3B6 | *Bradypus* | + | - | + | + | - | - | - | - | - | - | - | - | - | - | - |

SA = *S. aureus*; EC = *E. coli*; PA = *P. aeruginosa*; BS = *B. subtilis*; CA = *C. albicans*; SAg = *S. agalatiæ*; SS = *S. saprophyticus*; SE = *S. epidermidis*; KP = *K. pneumonie*; SM = *S. marcenses*; Cy = *Cytospora* sp.; Co = *Colleotrichum* sp.; Fu = *Fusarium* sp.; Hy = *Hypocrea* sp.; Tr = *Trichoderma* sp.

### Identification of bacterial isolates

For identification of the antibiotic-producing isolates, each one was grown individually (ISP2 agar, 1-4 weeks). Then, DNA was extracted according to the protocol described by Chun and Goodfellow (1995). DNA samples were quantified using a NanoDrop 2000 spectrophotometer (Thermo Fisher, USA). Polymerase chain reaction (PCR) protocols described by Cambronero-Heinrichs et al. (2019) to amplify 16S rRNA with primers 27F and 1492R were used. PCR products were visualized on agarose gels (1%) stained with Gel Red (Biotium Inc., USA). The PCR products were sequenced using BigDye Terminator v3.1 cycle sequencing kit and cleaned with BigDye XTerminator purification kit (Applied Biosystems, USA). Products were analyzed on a 3130xl Genetic Analyzer (Applied Biosystems, USA). Forward and reverse sequences were assembled using Bioedit. The obtained consensus sequences were compared against 16S rRNA bacterial/archaeal database of the NCBI using BlastN. All sequences were deposited on the Genbank under accessions OM638436-OM638444.

### Biosynthetic gene clusters (BGCs) analyses of antibiotic-producing isolates

For biosynthetic gene clusters (BCGs) analyses, the nine bioactive isolates were grown on ISP2 media (5 days at 30 ºC). Specimens classified within the same genera were combined in sterile microcentrifuge tubes with a difference in OD600 of 1:2 to achieve a good binning of the genomes according to their abundance. DNA was extracted using a DNA isolation kit (PowerSoil®, MoBio, Carlsbad, CA, USA) following the manufacturer’s instructions. DNA was quality checked and subjected to a 250 nt paired-end sequencing (Novoseq, Novogene). The raw sequences from this study were deposited in the GenBank database BioProject ID PRJNA804868

For the construction of the Metagenome Assembled Genomes (MAGs) sequences were quality filtered using bbmaptools (function bbduck) and then quality checked with FastQC. Sequences were assembled with MEGAHIT (Li et al., 2016). Only contigs larger than 1500 bp were further analyzed. The open reading frames were mapped using Prodigal v2.6.3 (Hyatt et al., 2010) and the single-copy core genes were identified with HMMER v3.3 (Johnson et al., 2010). The taxonomy of the SCG genes was estimated using DIAMOND (Buchfink et al., 2015). Manual binning was performed using Anvio v6.2. A bin was considered a MAG if it presented 100% completeness and >5% redundancy. Four bins were obtained from which two were classified as MAGs. Phylogenetic analysis of the four bins with closely related genomes (see table 2 for Genbank accessions) was performed by using 71 single-copy core genes (a curated list of genes available in Anvi’o phylogenomic workflow (https://merenlab.org/tutorials/infant-gut/#chapter-iii-phylogenomics) shared among the different genomes. Sequences were concatenated and further aligned using MUSCLE v3.8.1551 (Edgar, 2004). Phylogenetic relationship was estimated using FasTree v2.1 (Price et al., 2009).

**Table 2.**
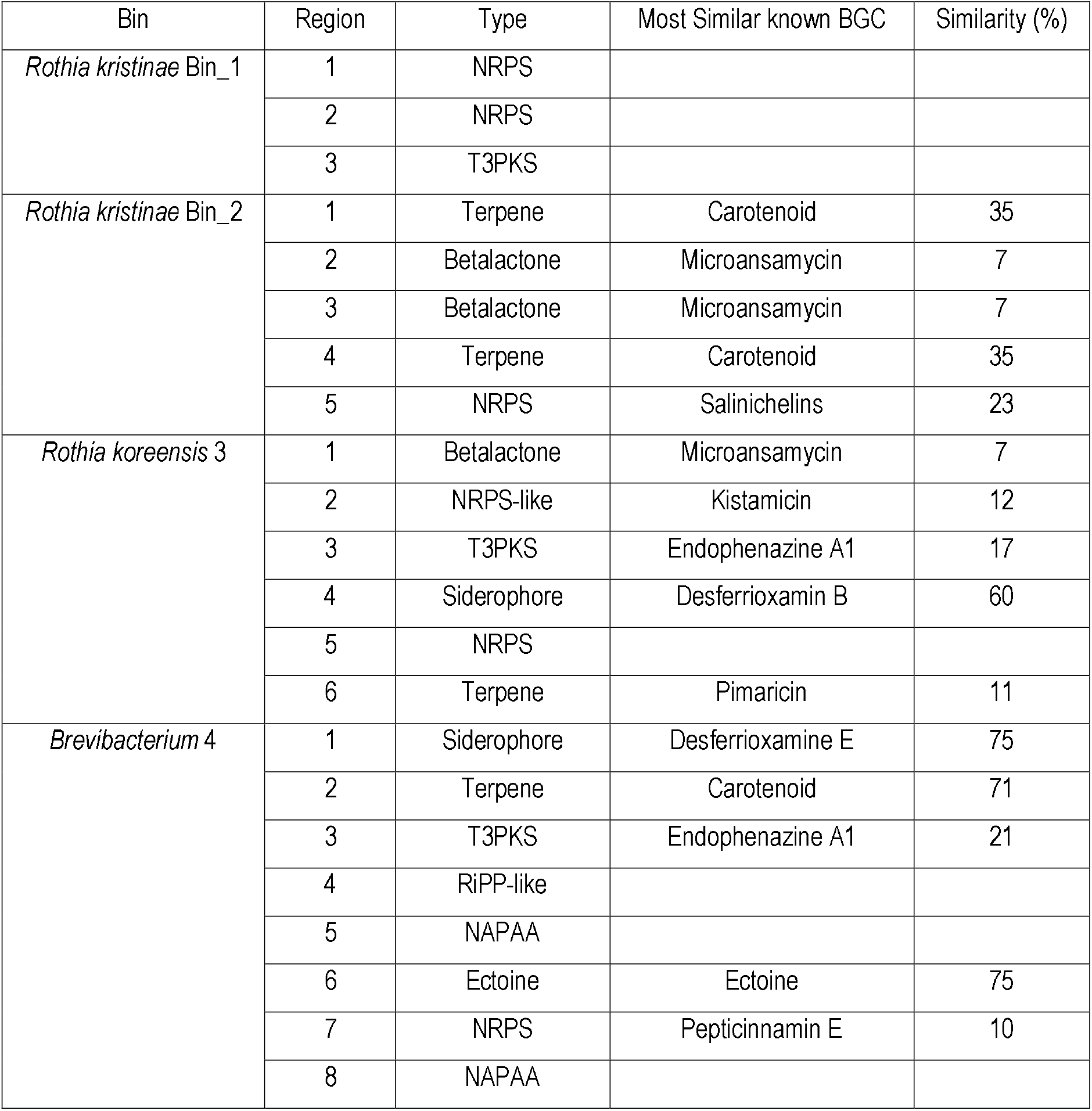
Putative biosynthetic gene clusters in *Rothia koreensis* (Bin 3), Rothia kristinae (Bin 1 and Bin 2) and *Brevibacterium* (Bin 4). BGCs were identified using Antismash 5.0 and classified agains the MiBiG database as described in “Experimental procedures”.

BGC analyses were conducted employing antiSMASH 5.0 (Blin et al., 2019) with default relaxed detection parameters that identify well-defined complete clusters and partial clusters missing one or more functional parts. We applied the following criteria for determining a known BGC: 100% of its genes have a significant BLAST hit to the corresponding genes of a known compound in the MIBiG database.

## Results and Discussion

### SEM reveals a high density of cocci-and bacilli-shaped microorganisms in sloths hair

Similar to the results obtained by Wujek and Cocuzza (1986), we observed through a scanning electron microscope (SEM) that the hair of sloths of the genus *Choloepus* has a series of longitudinal furrows and an average width of 138 ± 13 μm. On the other hand, the hair of sloths of the genus *Bradypus* is smoother, with irregular fissures throughout the entire hair and an average width of 261 ± 36 μm. Hair samples from both *Choloepus* and *Bradypus* (Fig. 2), showed a high density of cocci- and bacilli-shaped microorganisms. This high density of microorganisms was more noticeable in the hair of the sloths of the *Choloepus* species, which could be explained by the morphology. As observed in the SEM micrographs, the microbial communities are preferentially established within the furrows of the hair of the genus *Choloepus*. Presumably, this location can be favored by a lower exposure to mechanical stress, and greater availability of water and nutrients with respect to the ridges of the hair. Most of the microorganisms observed in those habitats correspond to coccoid bacteria, nevertheless, some hyphae-like structures and rod-shaped bacteria were present in less abundance. In *Bradypus* hair, a lower microbial density was observed and its location was not necessarily associated with the fissures throughout the entire hair. In *Bradypus*, it is also possible to observe coccoid bacteria, as on the hair of *Choloepus*.

**Figure 2.**
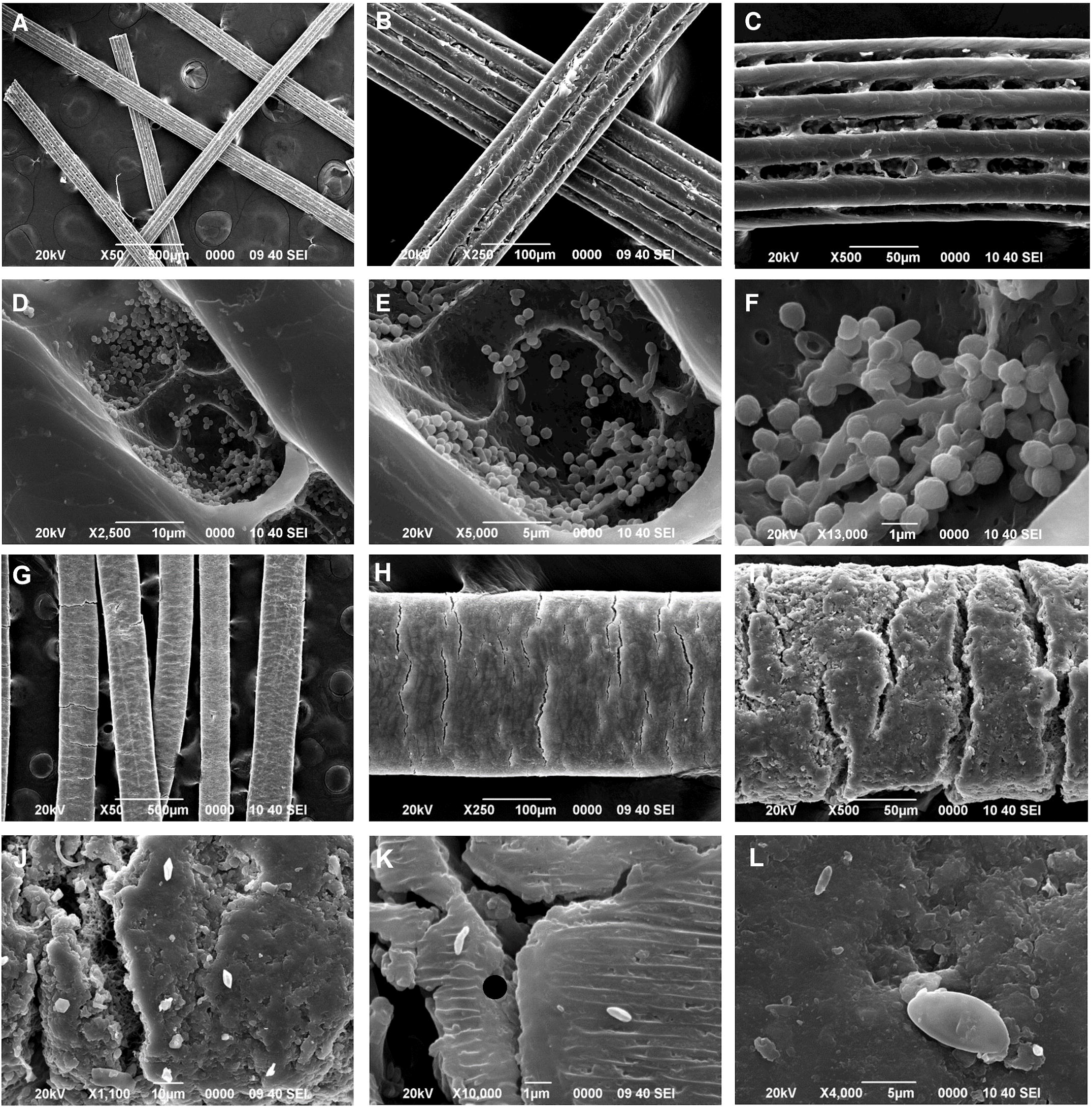
SEM images of the hair of Choloepus hoffmani and *Bradypus variegatus*. A) *Choloepus* hair are filament-like structures B) Filaments present a width of 138 ± 13 μm. C) The hair present multiple cavities along its longitude D) These cavities seem to be the niche for the microbial community inhabiting sloth hair. E) The higher abundance of microorganisms in these furrows is probably associated to the lack of mechanical stress they suffer F) Most microorganisms presented a coccoid-shape morphology, also some rod-shaped microorganisms were present. G) *Bradypus* hair is smoother H) It have average width of 261 ± 36 μm. I) It present irregular fissures throughout the entire hair longitude. J) Microorganisms survive in hair surface K) Coccoid and bacilli shaped bacteria are observed across the hair. L) The density of microorganisms is lower probably as a consequence of the hair morphology.

Since the sloths live in the refuge, the development of the characteristic greenish color in their fur was not observed, typically associated with the presence of the green algae *Trichophilus*. In the SEM photos, we were not able to determine the presence of cells of this algae either. Its absence is not rare, since the appearance of this algae occurs in wild sloths as a camouflage mechanism, which is not required in individuals living in captivity. Despite the absence of green algae *Trichophilus*, these first observations show that sloth hair is an environment rich in microorganisms. To reveal the identity of these microbial communities we decided to perform an analysis of 16S rRNA gene amplicons.

### Analysis of 16S rRNA gene amplicons indicates that sloth hair is governed by members of the Micrococcales order and shaped by the environment

The bacterial community inhabiting the hair of both sloth genera showed high diversity values. The average estimations of the Shannon-index were 3.06 and 2.72 for *C. hoffmani* and *B. variegatus*, respectively (see Figure S1). Wilcoxon test revealed a statistically significant difference (Shannon-index, p = 0.045) in the diversity of the microbial communities between the two sloths, in which *Choloepus* showed higher diversity than *Bradypus*; although multiple phylotypes were identified in all samples. In total, 4892 amplicon sequence variants (ASV) were identified in the 28 samples. The most abundant bacterial phyla were Actinobacteriota (10.82-96.44%), Firmicutes (1.60-65.56%), Proteobacteria (0.53-36.75%), and Bacteroidota (0.53-32.81%). Other phyla were detected in lower proportions, such as Acidobacteriota (0-1.91%), Chloroflexi (0.59-2.87%), Cyanobacteria (0-0.59%), Desulfobacteria (0-0.62%), Gemmatimonadota (0.84-3.46%) (see Figure 3A and Table S2).

**Figure 3.**
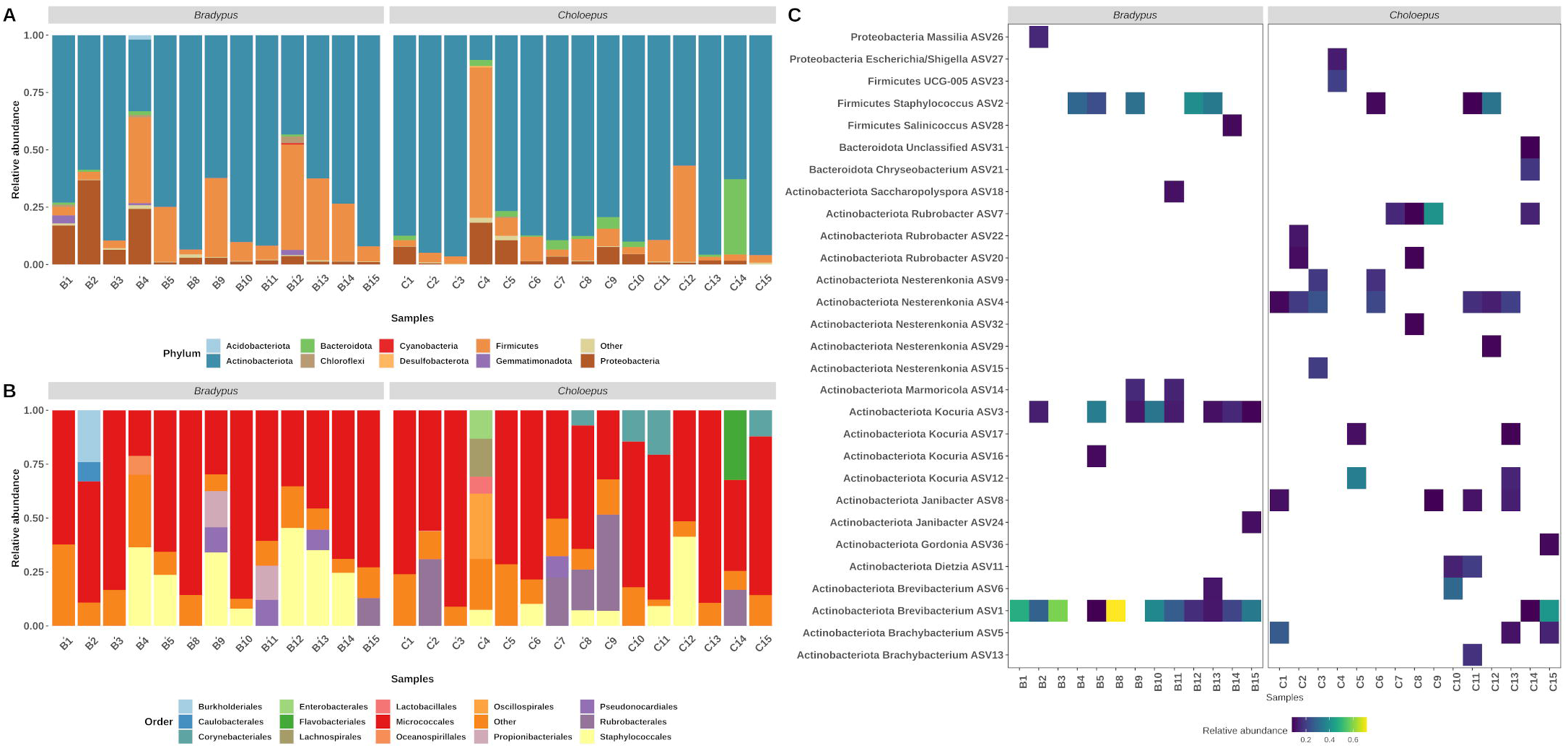
Taxonomic composition of prokaryotic community inhabiting the hair of *Bradypus variegatus* and *Choloepus hoffmani*. Relative abundance of bacterial and archaeal organisms at the A) phylum, B) Order levels. C) Relative percentage of 16S rRNA gene sequences assigned to each ASV (y axis) across the 28 samples analyzed (x axis).The ASV were taxonomically classified using SILVA reference database v138, as described in “Experimental procedures”. *Bradypus* samples are identified as B1 to B13 and *Choloepus* samples are identified as C1-C15.

At the order level, we observed that the bacterial community inhabiting the hair of both species of sloths was mainly dominated by members of the Micrococcales (see Figure 3B). Within this order, the families Brevibacteriaceae (0.70-75.30%) and Micrococcaceae (2.61-71.23%) were the most abundant (see Figure S2). Members of the Brevibacteriaceae family have been previously isolated from a variety of environments including dairy products (Forquin et al., 2009), poultry skin (Forquin-Gomez et al., 2014), insects (Katı et al., 2010), mural paintings (Heyrman et al., 2004), clinical samples (Roux and Raoult, 2009; Talento et al., 2009), saline sediments (Chen et al., 2016), mammal skin (Collins et al., 1983) and human microbiota (Forquin-Gomez et al., 2014). Most of the organisms found in this family are characterized as being halotolerant with the ability to produce sulfur volatile compounds (Arfi et al., 2003). Members of the Micrococcaceae family are usually associated with the skin microbiota of mammals (Collins et al., 2000), blood cultures (Hansen, 1985), and in various soil samples including marine sediments (Kim et al., 2004). These microorganisms have demonstrated a vigorous metabolism and an exceptional ability to adapt to harsh conditions (Dastager et al., 2014). Additionally, even though some members of both families ie. Brevibacteriaceae and Micrococcaceae have been associated with bacteremia and other infections (Asai et al., 2019; Bal et al., 2015; Kandi et al., 2016; Ramanan et al., 2014), most of the microorganisms belonging to this group are commensals frequently found on the skin and oral cavities of mammals (Collins et al., 1983; Dastager et al., 2014).

Furthermore, a significant proportion of members of the Staphylococcales (1.08-45.39%) and Rubrobacterales (0.17-44.57%) were found in the hair samples of *Bradypus* and *Choloepus* specimens (Figure 4B). Members of the family Staphylococcaceae are commonly found in the skin of multiple mammals (Lory et al., 2014). Particularly *Staphylococcus aureus* is a widely-known pathogen associated with multiple infections and has become more deadly due to the increase of antibiotic resistance (Hassoun et al., 2017). The prevalence of members of the Staphylococcaceae family in skin samples has been previously reported in multiple mammals, particularly in those mammals that have a strong anthropogenic influence (Ross et al., 2018) which is the case of the sloths sampled, that dwell in a human-controlled refugee. Contrarily to Staphylococcaceae, Rubrobacteriaceae members are characterized for being extremophilic bacteria, which have shown to survive extreme radiation, including gamma radiation (Ferreira et al., 1999). Members of this family are pink-pigmented bacteria commonly isolated from hot springs (Carreto et al., 1996), biodeteriorated monuments (Jurado et al., 2012), marine sponges (Kämpfer et al., 2014) and deep-sea sediments (Albuquerque et al., 2014). The presence of this taxon in sloth hair is probably associated with the high temperatures (the Sloth Sanctuary is located in Cahuita, with an annual temperature range of 21 to 31 ºC) and the constant exposure to UV radiation that the sloths endure in their niche (Duan et al., 2017; Gohli et al., 2019; Sun et al., 2018). Nevertheless, further study is required to comprehend the association between these extremophilic microorganisms and their hosts.

**Figure 4.**
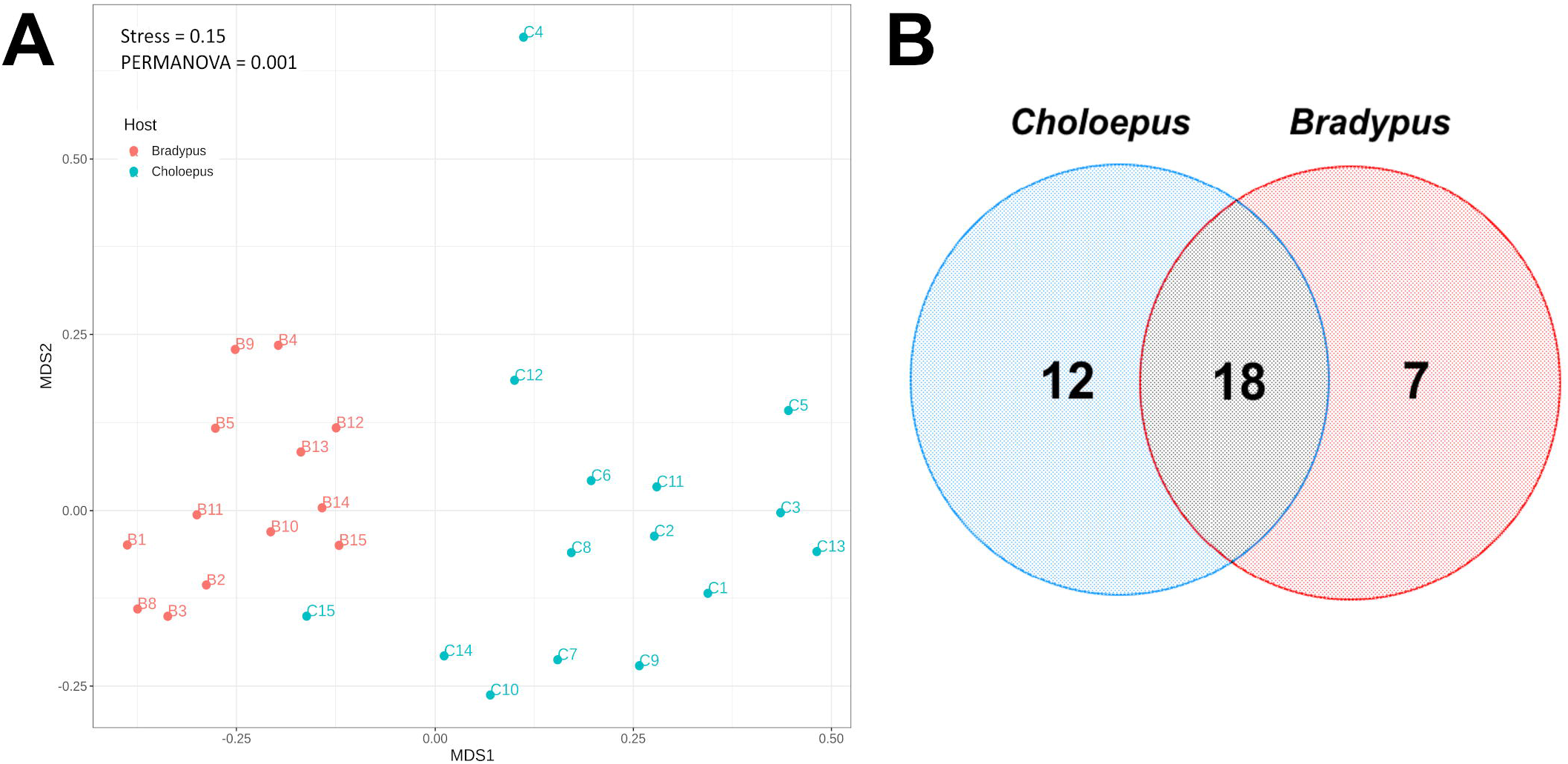
Non-metric multidimensional scaling (NMDS) analysis and Venn diagram of the prokaryotic communities in the hair of both sloth’s species. A) The NMDS analysis revealed a clustering of communities according to the host is shown. The NMDS and the permanova analyses were performed with Package Vegan v2.5-6. B) Venn diagram presenting the number of ASV of the core microbiome shared between *Bradypus variegatus* and *Choloepus hoffmani*. Both sloths share almost 50% of the core microbiome. Core microbiome was calculated using the R package microbiome as described in Experimental procedures.

As seen in Figure 3C, abundant ASVs detected in sloth’s hair encompass members of the genera *Brevibacterium (*0.70-75.30%), *Kocuria (*0.94-47.17%), *Staphylococcus* (1.01-45.09%), *Brachybacterium* (0.60-31.07%), *Nesterenkonia* (0.04-69.61%) and *Rubrobacter* (0.17-44.57%). These genera are characterized for inhabiting sediments usually with a high salt concentration (Chander et al., 2017; Dastager et al., 2014; Duan et al., 2017; Forquin-Gomez et al., 2014; Stackebrandt, 2014). The Sloth Sanctuary in Cahuita is located just 1.7 km from the Atlantic coast of Costa Rica, where high concentrations of salt in the environment are expected to occur. Therefore, it is not surprising that the microorganisms that inhabit the hair of the sloths in this refuge have a high content of bacteria associated with habitats with high concentrations of salt, thus being a factor that shapes the microbial community. It has also been shown that compounds secreted by the glands of the host can shape the microbial community through the variation of pH, salinity, and osmotic pressure (Byrd et al., 2018; Grice and Segre, 2011; Sanford and Gallo, 2013). The presence of microorganisms associated with saline sediments on our hair samples agrees with previous studies that showed that mammals’ skin microbiome resembles the community from the environment, including soils and sediments (Ross et al., 2018).

Specifically, sequences classified as *Brevibacterium* ASV1 and ASV6 were abundant ASVs particularly in *Bradypus* samples (see Figure 3C). *Brevibacterium* is a highly heterogeneous genus with multiple metabolic capabilities (Forquin-Gomez et al., 2014). It has been shown that this genus usually prefers to metabolize lactate as a source of carbon; lactate has been reported as a secreted compound by sebaceous mammals’ glands (Downey and Kealey, 2004). Additionally, this genus can also metabolize amino acids as carbon and nitrogen sources, probably allowing them to adapt well to the highly competitive hair niche on which algae, insects, protozoa, and other bacteria compete for resources (Mounier et al., 2007). Similarly, multiple sequences identified as *Kocuria* were found in samples belonging to both genera (*Choloepus* and *Bradypus*) (ASV3, ASV12, ASV16, and ASV17). *Kocuria* is characterized for mainly being aerobic chemoheterotroph microorganisms (Dastager et al., 2014). It usually expresses polymer degrading enzymes such as esterases (Kaur et al., 2011), and lipases (Reddy et al., 2003), suggesting that they might have the ability to degrade biopolymers found in the skin. A keratinase has also been found in *Kocuria* specimens (Bernald et al., 2006), which probably allow them to degrade the hair in which they inhabit to obtain energy.

Additionally, various abundant ASVs were classified as *Rubrobacter*. As mentioned above, the genus *Rubrobacter* is widely known for its ability to survive radiation and desiccation (Chen et al., 2004). Its resistance to these conditions probably allows members of this taxon to live on the sloth hair which is constantly exposed to UV radiation and desiccation due to the high sodium chloride content. Additionally, ASV classified as *Staphylococcus* were highly abundant, similar to what has been reported in human skin, where *Staphylococcus* and *Propionibacterium* are the main members of the community (Byrd et al., 2018; Cogen et al., 2008). It has been previously shown that *Staphyloccocus* is part of the normal hair microbiota in humans (Lousada et al., 2021). Even though some species such as *S. aureus* are important pathogens, most of the genotypes belonging to this group are commensal microorganisms that accomplish important processes for maturation of the immune system or secreted bacteriocins-like proteins that inhibit the proliferation of pathogenic bacteria (Nguyen et al., 2017; Newstead et al., 2020; Carson et al., 2017).

Finally, other sequence variants were assigned to the Actinobacteriota genera *Brachybacterium* (ASV5, ASV10, ASV13) and *Nesterenkonia* (ASV4, ASV9, ASV32, ASV29, ASV15). Both of these genera are poorly studied bacteria which, similar to *Brevibacterium* and *Kocuria*, show a great diversity in their niches, which range from plant growth promoters to human opportunistic pathogens (Edouard et al., 2014; Tak et al., 2018; Singh et al., 2016).

### The sampled two- and three-toed sloths share almost 50% of the core microbiome in their hair

Further Non-Metric Multidimensional and PERMANOVA analysis revealed that even though similar bacterial phyla were found in *Choloepus* and *Bradypus* hair, a significant difference between the bacterial communities was found (PERMANOVA, p = 0.001) (see Figure 4A). No significant difference was found between the sex (p = 0.18) or time in captivity (p = 0.143) for either species (data not shown). Previous analyses of the skin microbiome of multiple mammals found a specialization along mammal orders (Ross et al., 2018). It is reasonable to think that this specialization may also occur between sloth species. The difference in the microbial community observed could be associated with the physiology of each sloth as the compounds secreted by multiple glands in the skin would shape the microbial community.

Even though there is a significant difference in the microbial community between sloth species (see Figure 6A), our data revealed that the core microbiome of both sloths is similar (see Figure 4B). The core microbiome (according to the definition of Lahti and Shati (2017)) of both sloths consists mainly of members of the Actinobacteriota phylum and bacteria from the *Staphylococcus* genus, particularly of ASVs classified as *Brevibacterium, Staphylococcus, Kocuria, Nesterenkonia, Brachybacterium, Rubrobacter, Janibacter, Dietzia*, and *Saccharopolyspora*. The core microbiome of both sloth species in the Sanctuary shares almost 50% of the ASVs (see Figure 6B) which could suggest that these microorganisms exert a functional role in this habitat. Although the sampled sloths are in separate cages, not mixed between species, and the Sanctuary employees avoid handling the animals, the presence of a core microbiome that is shared between the two species must be validated with samples from other regions and with free-living animals. Despite this, our work provides a base and gives the first light of the microbiota that establishes itself in the fur of the sloths.

In an attempt to elucidate the role of the members of the microbiome in both sloth species and considering that many of these microorganisms belong to the phylum Actinobacteriota, we decided to evaluate the presence of bacteria that produce antibiotic molecules. Members of the phylum Actinobacteriota are known for their ability to produce antibiotic molecules, and many of them are found in association with higher organisms establishing symbiotic relationships (Matarrita-Carranza et al., 2021).

### Several members of the core microbiome are antibiotic-producing bacteria

To evaluate the capacity of the bacteria inhabiting sloth’s hair to produce antibiotic compounds, we isolated Actinobacteriota and evaluated their anti-microbial activity against multiple pathogenic microorganisms (see Experimental procedures). We were able to isolate 54 bacteria of which nine were antibiotic-producing. From these, five were isolated from *Choloepus* hair and four from *Bradypus* hair.

The antibiotic-producing bacteria were classified as either *Brevibacterium sediminis, Rothia koreensis*, or *Rothia kristinae*. Interestingly these genera were classified as part of the core microbiome in the microbial community analysis. *Rothia kristinae* and *Rothia koreensis* were only recently classified within the *Rothia* genus through phylogenomic analysis (Nouioui et al., 2018), before 2018 these bacteria were classified within the *Kocuria* genus according to its 16S rRNA sequence (Stackebrandt et al., 1995) suggesting that the absence of *Rothia* genus is a consequence of the close phylogenic relationship between *Rothia* and *Kocuria* that do not allow a clear separation through methodologies. Blastn of abundant ASV classified as Kocuria showed a 100% identity with *Rothia kristinae* or *Rothia koreensis*.

Biological activity was observed against *S. aureus, E. coli, P. aeruginosa, B. Subtilis. S. agalatiae* and *Cytospora* sp., however, the antimicrobial activity presented a wide variation even among strains of the same genus (see Table 1). Strains that showed activity against *S. aureus* (*R. koreensis* 10C1, *R. kristanae* 13C8, 9C1, 11C2, 3B2, and 3B6) showed no activity against *S. epidermidis* which is known to be a common symbiont in mammal’s skin (Liu et al., 2020). The ability of these isolates to inhibit well-known pathogens growing but not affect closely related symbionts may indicate that these Micrococcales may act as a protective mechanism principally against pathogens on sloth skin. Nevertheless, no inhibition was observed against other well-known pathogens such as *K. pneumonie, S. agalactiae*, and *S. saprohyticus*. Notably, *S. agalactiae* presented growth inhibition with isolate 3B2 an inhibition on its capacity to perform hemolysis when exposed to 9C1, 11C2, 3B2, and 13C9 (data not shown). Additionally, multiple strains presented activity against *Bacillus subtillis* which have been associated with food poisoning (13C8, 9C1, 11C2, 3B2, and 3B6).

Since many fungi have demonstrated the ability to infect the skin, we attempt to evaluate the antifungal activity of our isolates against fungi closely related to the ones previously found in *Bradypus* fur (Higginbotham et al., 2014) (see Experimental procedures). Three isolates (11C2,13C9, and 13C8) showed antibiosis against *Cytospora* sp. Despite *Cytospora* sp. is not a recognized mammal pathogen, its presence in sloth hair may indicate that these Microccocales have evolved to compete with other microorganisms that occupy a similar niche.

Furthermore, multiple isolates showed the ability to inhibit other Gram-negative bacteria such as *Pseudomonas aeruginosa* and *Escherichia coli* (see Table 1). *P. aeruginosa* is an opportunistic pathogen that has shown an intrinsic resistance to a variety of antibiotics (Gellatly et al., 2013; Qiu et al., 2009; Thi et al., 2020). Nowadays, *P. aeruginosa* strains resistant to almost all antibiotics (Molina-Mora et al., 2020) have been isolated and represent a significant health problem; therefore, isolates that show the ability to inhibit its growth are interesting for future research to identify the compounds responsible for this activity.

### BGCs found in antibiotic-producing bacteria are poorly studied but well conserved among strains of Micrococcales

When analyzing the Biosynthetic Gene Clusters (BGCs) in the assembled genomes, we were able to manually bin the sequences into four bins (see Table 2). Bins were classified as *Rothia koreensis* (Bin 3), *Rothia kristinae* (Bin 1 and Bin 2), and *Brevibacterium* (Bin 4). Furthermore, a comparison with closely related genomes obtained from the Genbank (see Figure 5) showed that our Bins have a very similar BGC composition in contrast with reference genomes of the same genus (see Figure 5) revealing that the BGCs are well conserved among multiple strains of the same genus.

**Figure 5.**
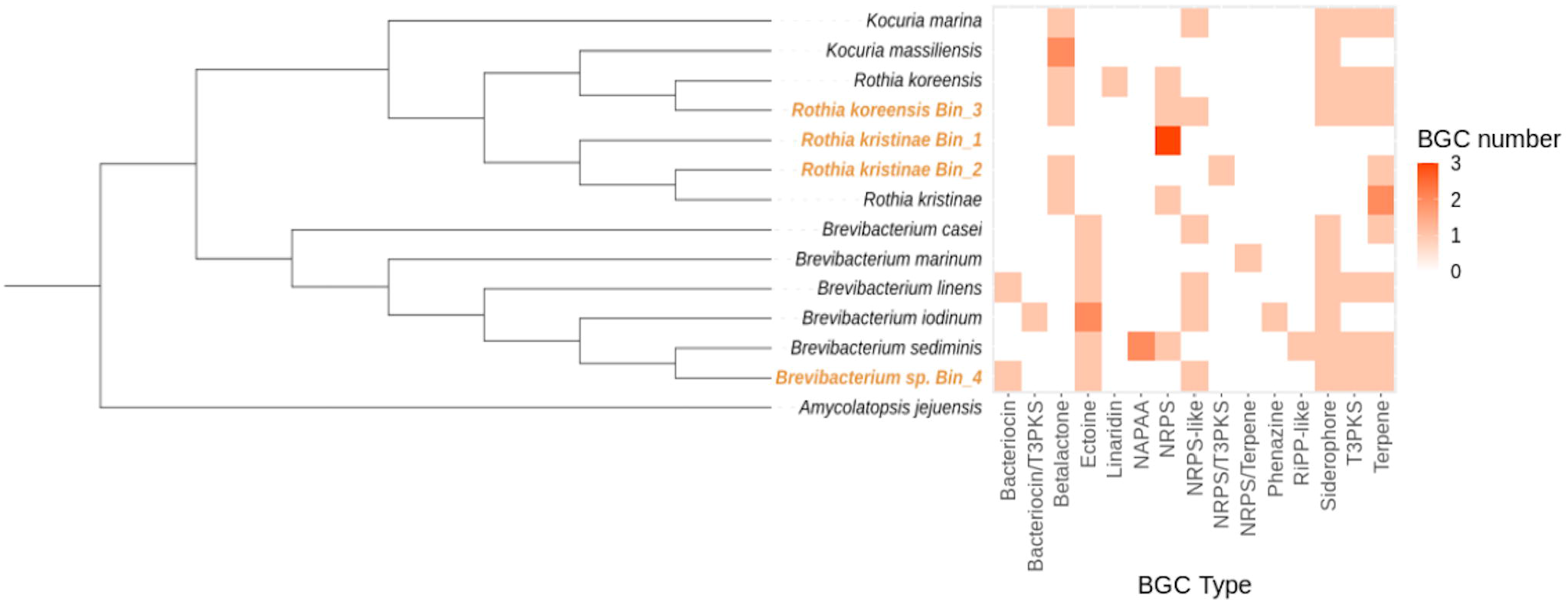
Heat map of BGCs types from the MAGs assembly and references genomes identified in antiSMASH and BiG-SCAPE.

Bin 4 is the MAG with more BGCS with a total of eight BGCS classified as siderophores, terpenes, T3PKS, RiPP-like, NAPAA, ectoine, and NRPS (Figure 5). However, none of the BGCs found in this genome showed a 100% similarity with any known cluster on the MiBiG database therefore, we cannot ensure that any specific metabolite is produced (Table 3). Nevertheless, BGC 22.1 was classified as a RIPP-like (see Table 2) and present a high similarity with Linocin M18 (81%), suggesting that a similar bacteriocin may be produced. Linoncin M18 is a bacteriocin produced by *Brevibacterium linens*; this peptide has been shown to be active against Gram-positive bacteria, particularly *Bacillus subtilis, Staphylococcus aureus*, and *Listeria monocytogenes* (Valdés-Stauber et al., 1994). Furthermore, other bacteriocins have been shown to be responsible for the antibiotic activity of different *Brevibacterium* strains; specifically Linencin A (Forquin-Gomez et al., 2014) and Linenscin OC2 (Maisnier-Patin et al., 1995). Linencin A is a 95 KDa peptide that inhibits the growth of only other *B. linus* strains. On the other hand, Linenscin OC2 showed to inhibit the growth of *Arthrobacter, Microccus*, and *Corynebacterium*.

Furthermore, Bin 1 and Bin 2 classified as *Rothia kristinae* showed the presence of NRPS (three BGCs), terpenes (two BGCs), beta lactone (two BGCs), and a T3PKS. Finally, Bin 3 showed six BGCs classified as terpene (one BGCs), beta lactone (one BGCs), NRPS-like (one BGCs), NRPS (one BGCs), T3PKS (one BGCs), and a siderophore (one BGCs). No BGCs found in the MAGs present a 100% similarity with the ones reported on the MIBIG database, therefore we can not ensure completely that any cluster produces a known secondary metabolite molecule. The low identity of the clusters found in these bins may indicate that novel metabolites are being produced, ensuring the potential of these microorganisms for the bioprospecting of novel antibiotics. Members of the *Rothia* genus have been poorly described as antibiotic-producing bacteria with only a few reports found (Uranga et al., 2020). Particularly, a porcine commensal classified as *Rothia nasimurium* showed the genetic capacity to produce the ionophore antibiotic valinomycin (Roiser et al 2017), but no identification of the compound was performed.

In summary, this work characterizes the bacterial community inhabiting the hair of *Choloepus hoffmani* and *Bradypus variegatus* in a group of animals in captivity. Analysis of the V4 region of the 16S rRNA gene showed that Actinobacteriota, Proteobacteria, and Firmicutes were the most dominant phyla, and *Brevibacterium, Janibacter, Rubrobacter, Brachybacterium, Kocuria*, and *Staphylococcus* the most dominant genera across all samples. Additionally, further analysis revealed that even though a significant difference exists between the community inhabiting both sloths’ sloths’ species, the phylotypes that conform to the core microbiome were mostly shared. Isolation of bacteria revealed that phylotypes closely related to the core microbiome exert an antibiotic activity against a wide variety of pathogens. Furthermore, metagenomics made from the antibiotic-producing strains revealed that multiple BGCs were found in these isolates including siderophores, terpenes, betalactone, Type III polyketide synthases (T3PKS), ribosomally synthesized and post-translationally modified peptides (RiPPs) similar to bacteriocin Linocin M18, non-alpha poly-amino acids (NAPAA) like e-Polylysin, ectoine and nonribosomal peptides (NRPs).

Our results suggest the presence of antibiotic-producing bacteria in the fur of sloths that could modulate the microbial community in that ecosystem. This modulation makes it possible to control the proliferation of potentially pathogenic bacteria for the host or inhibit other competitors (e.g. fungi) in the niche. On the other hand, our results point to sloth fur as a new source of antibiotic-producing bacteria that may open the opportunity for the discovery of new bioactive molecules.

## Acknowledgements

We thank Sofía Vieto, Roberto Avendaño and Felipe Vasquez-Castro for their help in collecting samples or processing them in the laboratory.

## Author Contribution

MC and DRG conceived and designed the experiments; DRG, KVM, RP performed the experiments; DRG, MC and KRJ analyzed the data; MC, JA-A and RP contributed reagents or materials; DRG and MC wrote the paper. All authors reviewed and approved the final version of the manuscript.

