## Supplementary information for "Antibiotic-producing Micrococcales govern the microbiome that inhabits the fur of two- and three-toed sloths"

^1^Centro Nacional de Innovaciones Biotecnológicas (CENIBiot), CeNAT-CONARE, 1174-1200, San José, Costa Rica, ^2^Escuela de Biología, Universidad de Costa Rica, 11501-2060, San José, Costa Rica; ^3^Laboratorio Nacional de Nanotecnología (LANOTEC), CeNAT-CONARE, 1174-1200 San José, Costa Rica. ^4^The Sloth Sanctuary of Costa Rica, Limon, Costa Rica, ^5^Escuela de Química, Universidad de Costa Rica, 11501-2060, San José, Costa Rica, ^6^Centro de Investigaciones en Productos Naturales (CIPRONA), Universidad de Costa Rica, 11501-2060, San José, Costa Rica.

Keywords: Sloths, Micrococcales, Antibiotic-producing bacteria, *Kocuria, Brevibacterium*, *Rothia*, Hair microbiota

*** Correspondence to: Max Chavarría**

Escuela de Química & Centro de Investigaciones en Productos Naturales (CIPRONA)

Universidad de Costa Rica

Sede Central, San Pedro de Montes de Oca, San José, 11501-2060, Costa Rica

Phone (+506) 2511 8520.  Fax (+506) 2253 5020

**LEGENDS OF SUPPLEMENTARY TABLE**

**Table S1. Metadata of sloths sample on the sloth sanctuary.**

**Table S2. DNA sequence and phylogenetic assignment of the most abundant ASVs detected in the hair samples using Illumina-based amplicon deep-sequencing.**

See Excel file.

**LEGENDS OF SUPPLEMENTARY FIGURES**

**Figure S1. Diversity measures of the hair samples from *Bradypus variegatus* and *Choloepus hoffmani***. The diversity measures (Shannon, Simpson and Observed Richness) were calculated using phyloseq. Figure shows A) diversity measures of all samples grouped by sample point. B) diversity measures of all samples.

**Figure S2. Taxonomic composition at the family level of prokaryotic community inhabiting the hair of *Bradypus variegatus* and *Choloepus hoffmani*.** Relative abundance of bacterial and archaeal organisms at the family level. The ASV were taxonomically classified using SILVA reference database v138 (Quast et al., 2013) as described in “Experimental procedures.” *Bradypus* samples are identified as B1 to B13 and *Choloepus* samples are identified as C1-C15.

**Figure S3. Isolates that present antimicrobial activity found in the hair of *Bradypus variegatus* and *Choloepus hoffmani.*** Bacteria were grown on ISP2 agar for 1 week. All bacteria are classified either as *Brevibacterium*, *Kocuria* or *Rothia* according to its 16S rRNA sequence.
