## Supplementary figures and images for "Antibiotic-producing Micrococcales govern the microbiome that inhabits the fur of two- and three-toed sloths"

### Supp. Fig S2

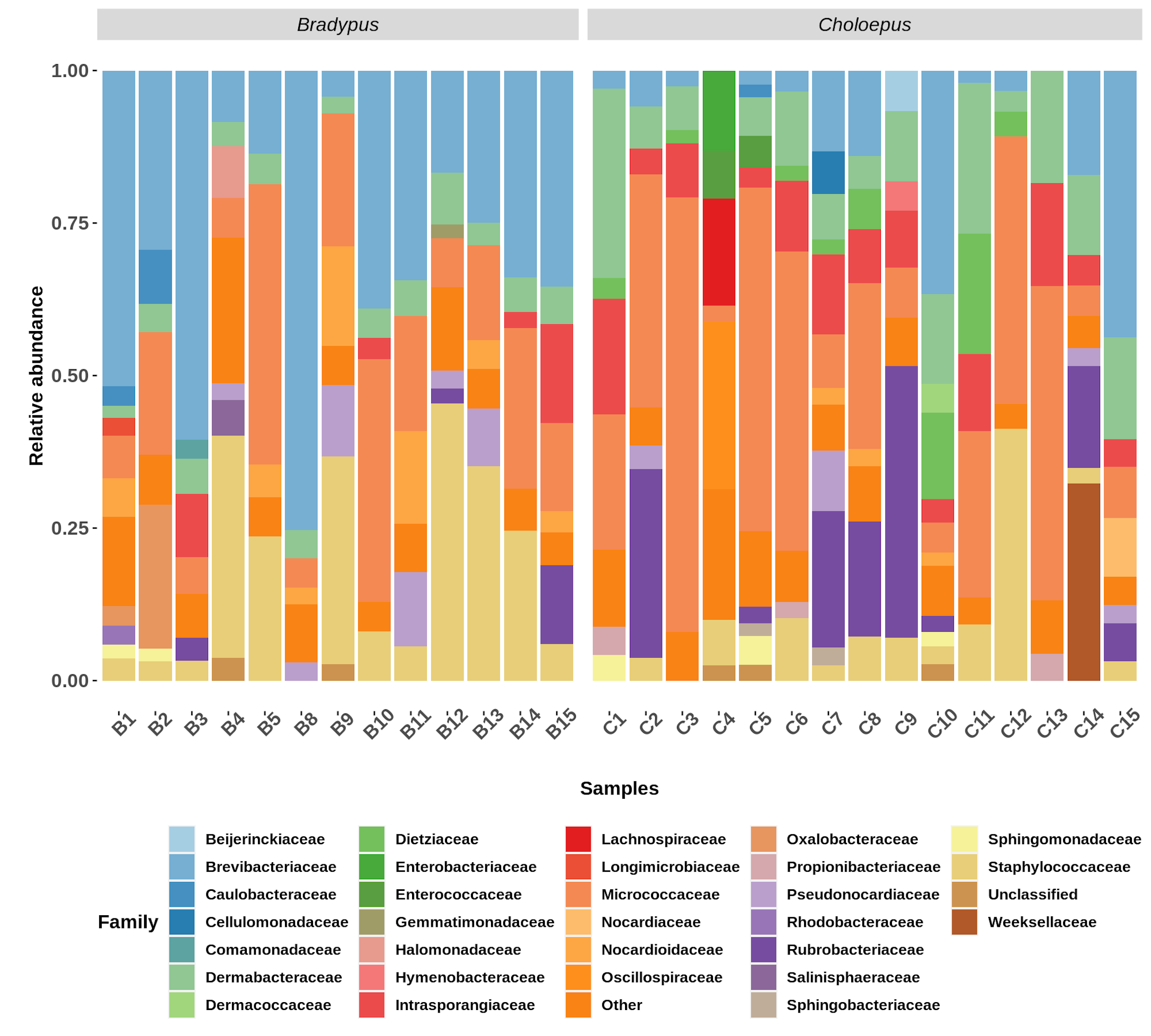

### Supp. Fig. S1

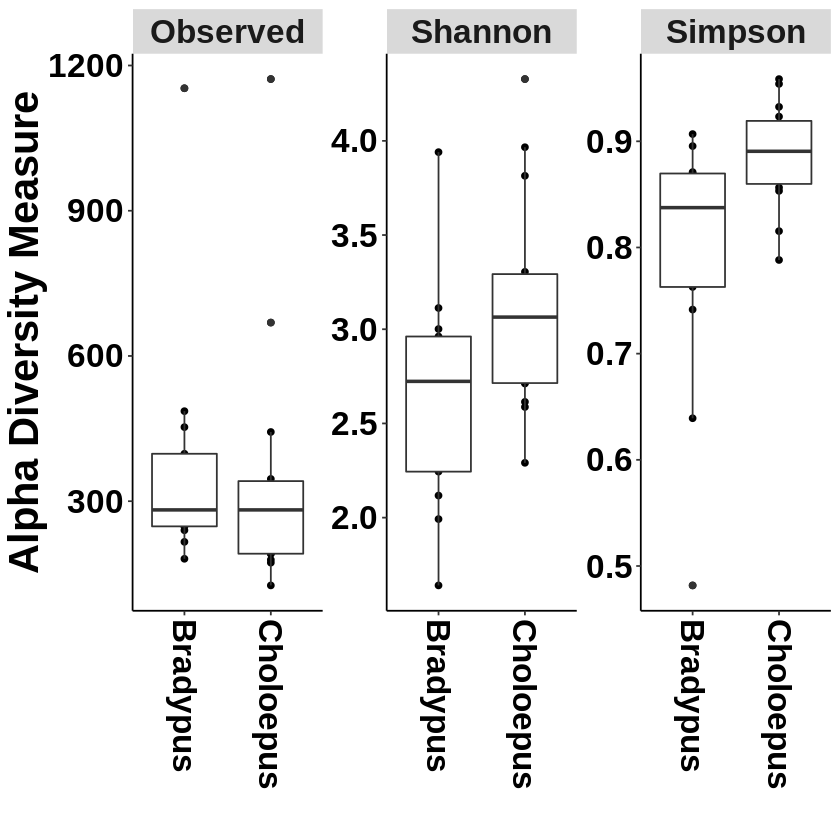

### Supp. Fig. S3

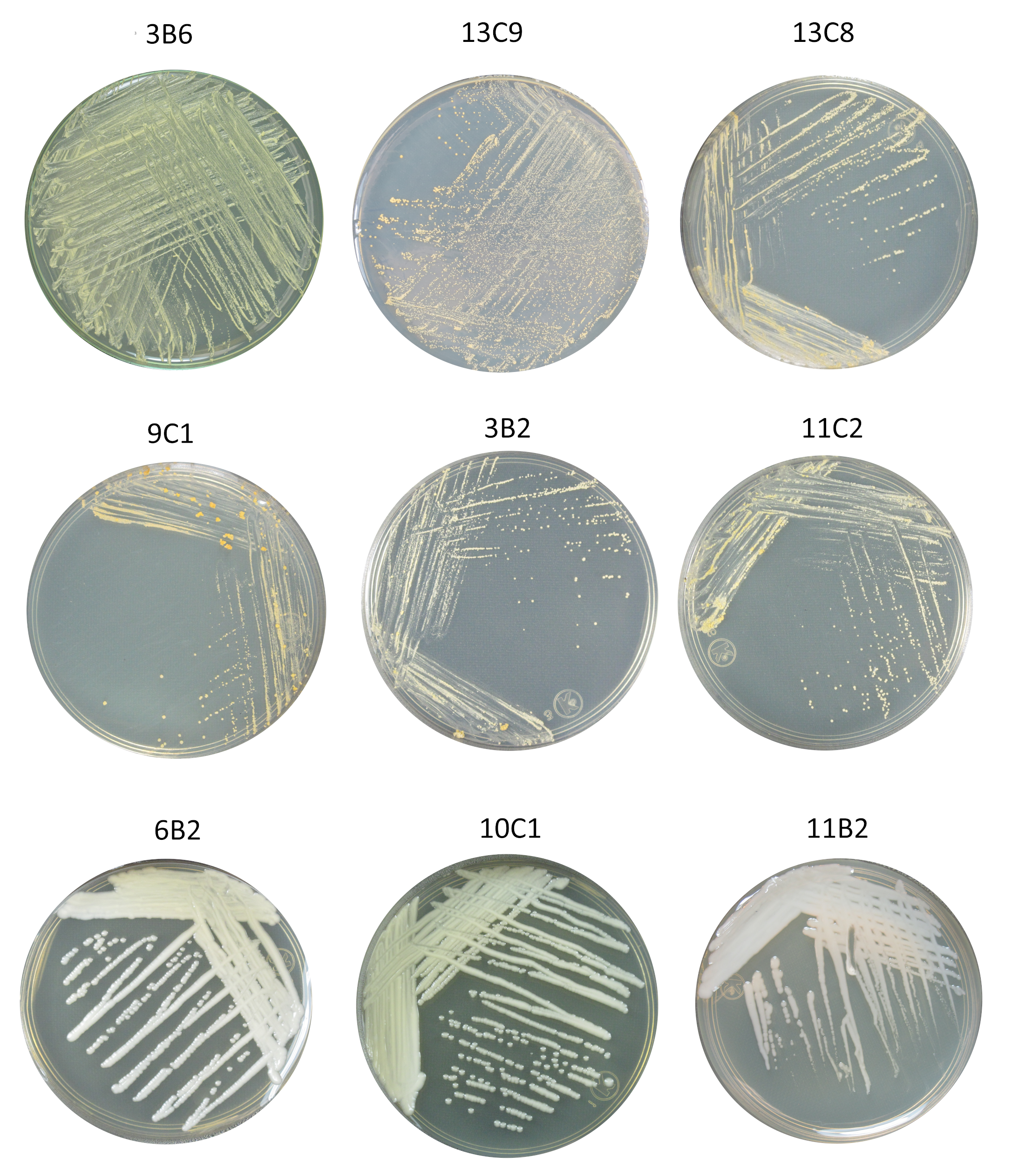
